# Human cell-dependent, directional, time-dependent changes in the mono- and oligonucleotide compositions of SARS-CoV-2 genomes

**DOI:** 10.1101/2021.01.05.425508

**Authors:** Yuki Iwasaki, Takashi Abe, Toshimichi Ikemura

## Abstract

**Background:** When a virus that has grown in a nonhuman host starts an epidemic in the human population, human cells may not provide growth conditions ideal for the virus. Therefore, the invasion of severe acute respiratory syndrome coronavirus-2 (SARS-CoV-2), which is usually prevalent in the bat population, into the human population is thought to have necessitated changes in the viral genome for efficient growth in the new environment. In the present study, to understand host-dependent changes in coronavirus genomes, we focused on the mono- and oligonucleotide compositions of SARS-CoV-2 genomes and investigated how these compositions changed time-dependently in the human cellular environment. We also compared the oligonucleotide compositions of SARS-CoV-2 and other coronaviruses prevalent in humans or bats to investigate the causes of changes in the host environment.

**Results:** Time-series analyses of changes in the nucleotide compositions of SARS-CoV-2 genomes revealed a group of mono- and oligonucleotides whose compositions changed in a common direction for all clades, even though viruses belonging to different clades should evolve independently. Interestingly, the compositions of these oligonucleotides changed towards those of coronaviruses that have been prevalent in humans for a long period and away from those of bat coronaviruses.

**Conclusions:** Clade-independent, time-dependent changes are thought to have biological significance and should relate to viral adaptation to a new host environment, providing important clues for understanding viral host adaptation mechanisms.

## Background

Severe acute respiratory syndrome coronavirus-2 (SARS-CoV-2), an RNA virus belonging to the betacoronavirus genus, began to spread in the human population in 2019. This viral strain is believed to have been originally prevalent in bats and transferred to the human population through intermediate hosts [1]. Viral growth requires a wide variety of host factors (nucleotide pools, proteins, RNA, etc.) and should evade the diverse antiviral mechanisms of host cells (antibodies, killer T cells, interferon, RNA interference, etc.) [2–4]. Since ancestral SARS-CoV-2 strains are thought to be endemic in bats, they should be well adapted to their host environment; when the virus invades the human population, human cells may not provide growth conditions ideal for the virus. For efficient growth and rapid spread of the infection, changes in the viral genome should be required. Analyses of time-dependent changes in SARS-CoV-2 in the human population can be used to characterize how and why viral genomes change to adapt to a new host environment.

Due to the great threat of COVID-19 and remarkable development of sequencing technology, a massive number of SARS-CoV-2 genome sequences are available in databases, even though the epidemic has lasted for approximately 10 months. These sequence data have provided a wide range of insights into SARS-CoV-2 [5,6]. Phylogenetic methods based on sequence alignment have been widely used in molecular evolution studies [7,8], and these methods are well refined and essential for studying phylogenetic relationships between different viral species and variations in the same viral species at the single-nucleotide level. However, when dealing with a massive number of genome sequences, methods based on sequence alignment become problematic because they require a large amount of computational resources.

We have continued to develop sequence alignment-free methods focused on the oligonucleotide compositions of genome sequences [9–12]. Notably, oligonucleotide composition varies widely among species, including viruses, and is designated as genome signatures [13]. These compositions can be treated as numerical data, and a massive amount of sequence data can easily be subjected to various statistical analyses. Furthermore, even genomic fragments without orthologous and/or paralogous pairs can be compared [11,12,14–17]. Specifically, our previous work on influenza A-type virus genomes found that the oligonucleotide compositions of the viral genomes differed between hosts (e.g., humans and birds), even for viruses within the same subtype (e.g., H1N1 and H3N2 of type A) [11,12,14]; we also examined changes in the oligonucleotide compositions of influenza H1N1/09, which have been epidemic in humans beginning in 2009, and found that their compositions changed to approach those of the seasonal flu strains H1N1 and H3N2 [11]. Furthermore, although epidemics of the H1N1 and H3N2 strains began several decades apart, these strains showed highly similar chronological changes from the start of these epidemics. These evolutionary yet reproducible changes suggest that mutations to adapt to a new host environment inevitably accumulate when the host species of a virus changes, and these changes can be efficiently detected by analyzing oligonucleotide compositions.

Several groups, including ours, have examined changes in SARS-CoV-2 genomes during the early stages of the SARS-CoV-2 epidemic and found clear directional changes in a group of mono- and oligonucleotides detectable on even a monthly basis [15,18,19]. These directional changes will allow us to predict changes in the near future. Notably, near-future prediction and verification should be the most direct ways to test the reliability of the obtained results, models and ideas (e.g., those discovered for influenza viruses), providing a new paradigm for molecular evolutionary studies. In this context, the present study analyzed the genome sequences of over seventy thousand SARS-CoV-2 strains isolated from December 2019 to September 2020.

## Results

### Directional changes in the mononucleotide compositions (%) of SARS-CoV-2

For fast-evolving RNA viruses, diversity within the viral population arises rapidly as the epidemic progresses and subpopulation structure forms; the GISAID consortium has defined at least seven main clades (G, GH, GR, L, V, S and Others). Notably, the elementary processes of molecular evolution are based on random mutations, and strains belonging to different clades are thought to have evolved independently. Therefore, the observation of highly similar time-dependent changes independent of clade has certain biological meanings and may be inevitable for efficient growth in human cells. From this perspective, we first examined time-dependent changes in the mononucleotide compositions (%) of SARS-CoV-2 strains isolated from December 2019 to September 2020.

Among the seven clades (G, GH, GR, L, V, S and Others) reported by the GISAID consortium, we used six clades (G, GH, GR, L, V and S), excluding Others, in the analysis. For the time-series analysis, we calculated the average mononucleotide compositions (%) of the genomes in each clade collected monthly; in Fig. 1A, the mononucleotide composition of each clade is shown as a colored line, while that for the monthly collected genomes belonging to all clades is shown as a dashed line.

**Fig. 1.**
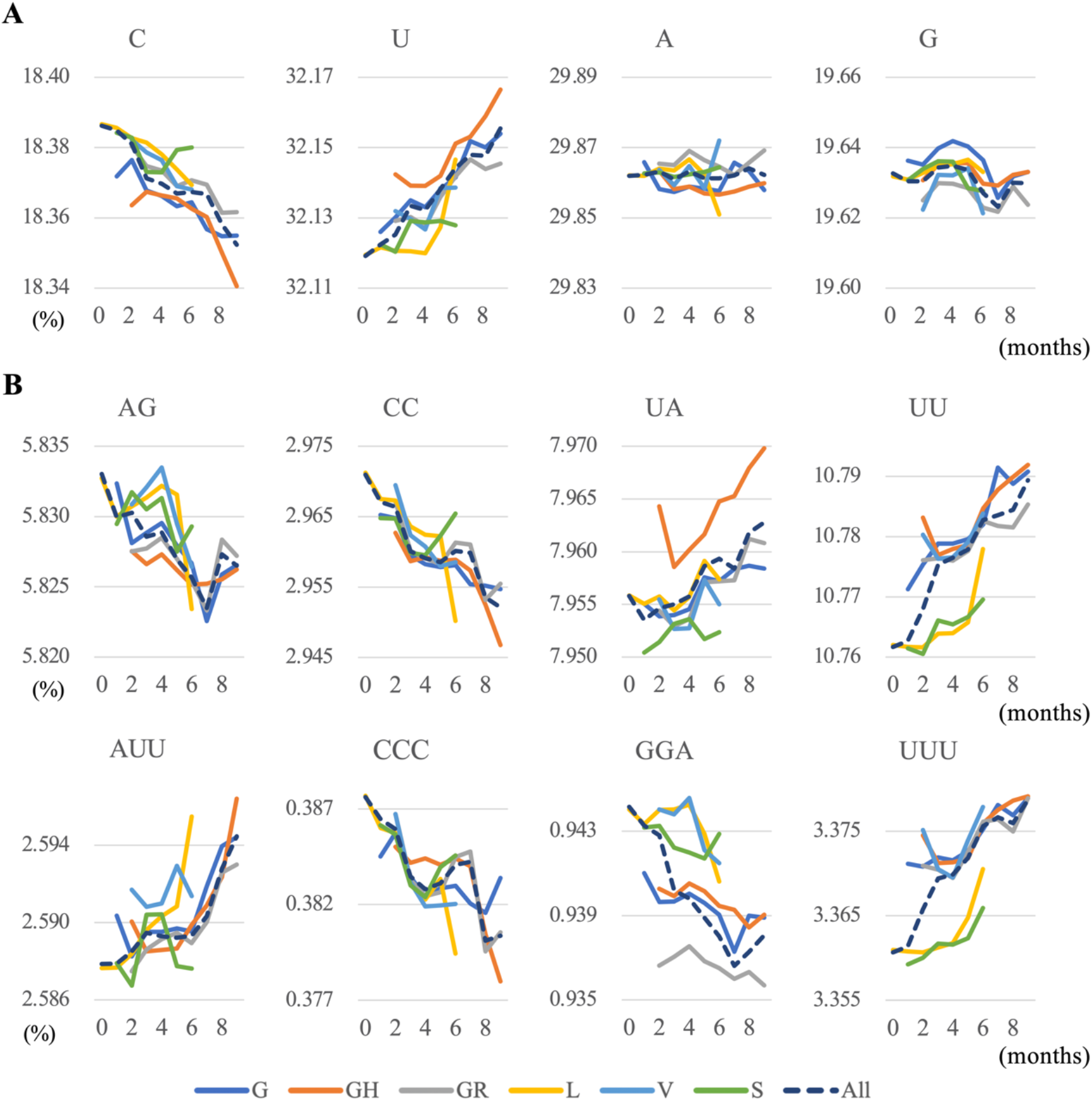
Time-dependent directional changes in nucleotide compositions. (A) Average mononucleotide compositions (%) in the SARS-CoV-2 genomes of each clade isolated in each month are plotted against the elapsed month. To compare the four mononucleotides, the scale widths on the vertical axis are set to the same values. The colored lines distinguishing the clade (G, GH, GR, L, V and S) are shown at the bottom of the figure. The dashed line shows the averaged compositions for all strains isolated in each month. (B) The average di- and trinucleotide compositions that primarily undergo common directional changes among the six clades are plotted against the elapsed month.

Regardless of clade, the composition of C decreased, while that of U increased in a time-dependent manner, but the changes in A and G composition were less clear (Fig. 1A). Correlation coefficients between the mononucleotide composition and month from the start of the epidemic showed a high negative correlation for C and a high positive correlation for U for all clades, but there was no clear directionality for A and G (Fig. 1A and Tables 1, 2). These results indicate that the mononucleotide composition of this virus may be prone to biased mutations that reduce C and increase U or the mutated strains tend to be more favorable for growth in human cells.

**Table 1.** Correlation coefficients for time-dependent changes in mono- and oligonucleotide compositions in SARS-CoV-2 that have increased.

|  | Clade G | Clade GH | Clade GR | Clade L | Clade V | Clade S |
| --- | --- | --- | --- | --- | --- | --- |
| U | 0.97 | 0.92 | 0.91 | 0.72 | 0.66 | 0.73 |
| UA | 0.89 | 0.80 | 0.92 | 0.62 | 0.30 | 0.50 |
| AUU | 0.81 | 0.78 | 0.92 | 0.90 | 0.27 | 0.05 |
| CAU | 0.56 | 0.59 | 0.54 | 0.51 | 0.55 | 0.17 |
| UGU | 0.68 | 0.82 | 0.52 | 0.59 | 0.88 | 0.50 |
| UUA | 0.96 | 0.72 | 0.93 | 0.80 | 0.19 | 0.34 |
| UUG | 0.67 | 0.78 | 0.20 | 0.92 | 0.69 | 0.05 |
| UUU | 0.94 | 0.82 | 0.89 | 0.80 | 0.41 | 0.93 |

**Table 2.** Correlation coefficients for time-dependent changes in mono- and oligonucleotide compositions in SARS-CoV-2 that have decreased.

|  | Clade G | Clade GH | Clade GR | Clade L | Clade V | Clade S |
| --- | --- | --- | --- | --- | --- | --- |
| C | -0.95 | -0.83 | -0.95 | -0.98 | -0.97 | -0.35 |
| AG | -0.77 | -0.71 | -0.27 | -0.57 | -0.67 | -0.44 |
| CA | -0.73 | -0.93 | -0.85 | -0.16 | -0.45 | -0.82 |
| CC | -0.93 | -0.87 | -0.75 | -0.90 | -0.90 | -0.09 |
| CU | -0.81 | -0.40 | -0.79 | -0.52 | -0.15 | -0.10 |
| GA | -0.28 | -0.90 | -0.80 | -0.62 | -0.73 | -0.29 |
| GG | -0.60 | -0.79 | -0.65 | -0.03 | -0.10 | -0.23 |
| UC | -0.33 | -0.23 | -0.10 | -0.58 | -0.93 | -0.41 |
| AGC | -0.57 | -0.87 | -0.41 | -0.80 | -0.78 | -0.12 |
| CCC | -0.69 | -0.81 | -0.67 | -0.94 | -0.81 | -0.50 |
| GAC | -0.73 | -0.91 | -0.65 | -0.35 | -0.18 | -0.55 |
| GAG | -0.11 | -0.58 | -0.64 | -0.50 | -0.81 | -0.02 |
| GGA | -0.73 | -0.84 | -0.75 | -0.65 | -0.81 | -0.52 |

### Directional changes in short oligonucleotide compositions

Oligonucleotides are known to act as functional motifs, such as binding sites for a wide variety of proteins and target sites for RNA modifications. Therefore, directional changes in some oligonucleotides independent of clade may relate to certain processes for adaptation to the new host environment. Our previous work on influenza A viruses found that their oligonucleotide compositions varied among prevalent hosts [11,12]; notably, although influenza virus isolated from humans tended to prefer A and U (but not G and C) more than viruses isolated from birds, the human viruses showed a preference for GGCG and GGGG, which are G- or C-rich. Importantly, there are various examples of oligonucleotides whose changes in composition cannot be explained by changes in mononucleotide composition alone, and these changes may relate to the molecular mechanisms of viral adaptation to a new host.

From this perspective, we next analyzed time-dependent changes in di- and trinucleotide compositions and found that a group of di- and trinucleotides showed a highly positive or negative correlation (Figs. 1B, S1 and Tables 1, 2). Interestingly, a group of A- or G-rich oligonucleotides, such as GAG and GGA, showed a high positive correlation independent of clade, which was not expected from the changes in mononucleotide compositions alone. To confirm the extent of these changes, we also calculated the fold change in composition for the first isolated month and the last examined month (Fig. 2) and found clear increases and decreases in mono- and oligonucleotide compositions common among the six clades, which supports the result presented in Fig. 1 and Tables 1 and 2.

**Fig. 2.**
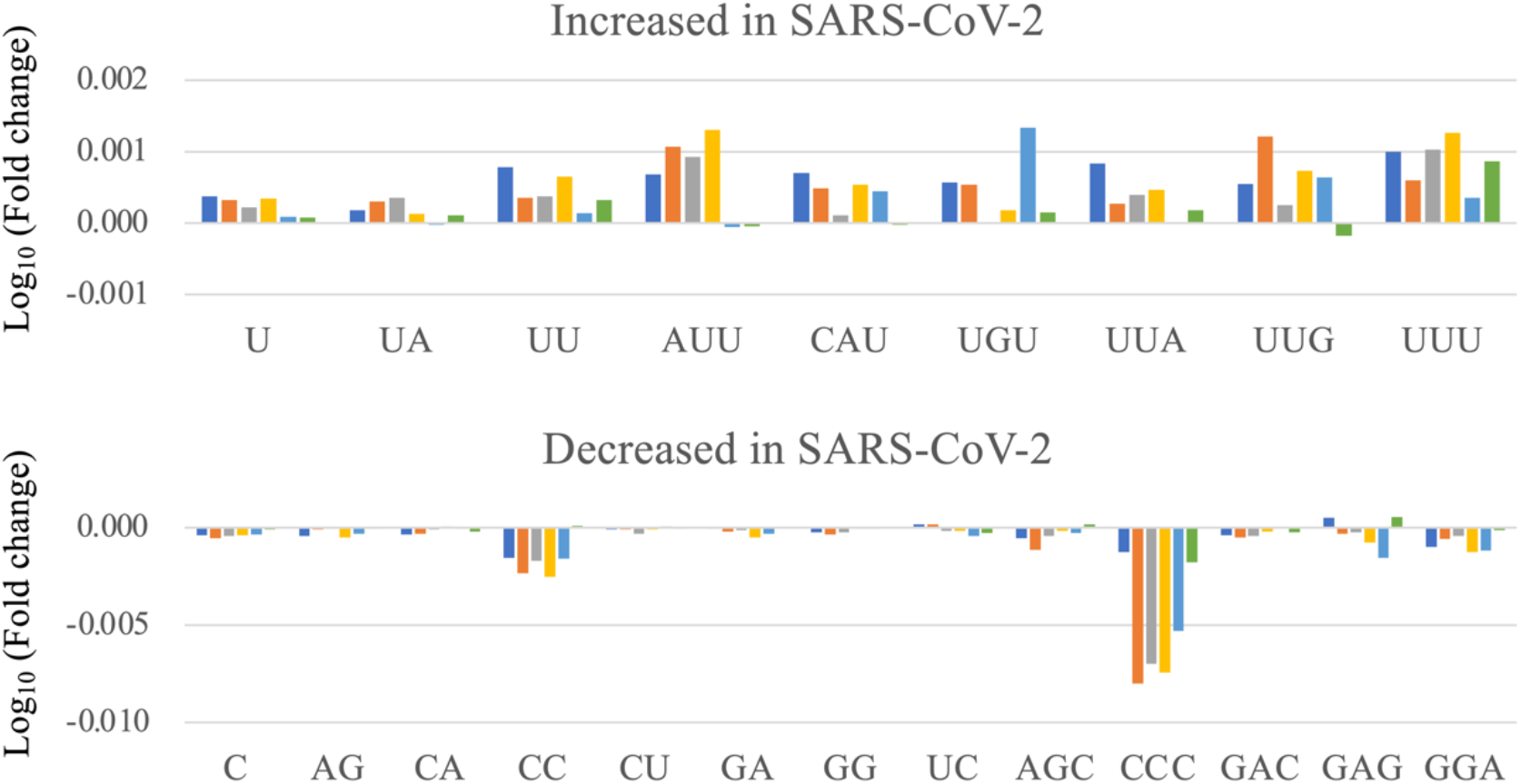
Fold changes in nucleotide composition between the epidemic start and the last month of analysis. A bar plot shows the fold change in composition of each mono- or oligonucleotide; this value was calculated by dividing the nucleotide composition in the last month of analysis by that at the start of the epidemic. Each bar is colored to indicate the clade, as described in Fig. 1. Since we analyzed strains belonging to different clades separately, data from the first or last month differed among clades; see also the Methods section.

### Changes towards the sequences of other coronaviruses prevalent in humans

In a previous study of SARS-CoV-2 [16], we analyzed mono- and dinucleotide compositions for the first four epidemic months without separating the sequences by clade. Notably, the directional changes shown in Figs. 1 and 2 and Tables 1 and 2 were absolutely consistent with the previous results, even when the six clades were separately analyzed. In the previous study, time-series analysis of ebolavirus at the beginning of the epidemic in West Africa in 2014 also showed directional changes in a group of mono- and dinucleotide compositions, but these directional increases/decreases tended to slow approximately 10 months after the start of the epidemic. The increase/decrease trend for SARS-CoV-2 is far from slowing after 10 months, and the next important questions are how long these directional changes in this virus will last and whether there are possible goals to these changes.

To conduct this near-future prediction, the following information concerning influenza viruses should be useful. As mentioned before, mono- and oligonucleotide compositions in influenza H1N1/09 changed towards those of seasonal influenza strains such as the H1N1 and H3N2 subtypes [11]. Furthermore, all the human subtypes showed directional changes away from the compositions of all avian influenza A subtypes and closer to those of the human influenza B type, which has been prevalent only in humans [14]. If we assume that changes similar to those in the influenza virus will occur, the mono- and oligonucleotide compositions of interest for SARS-CoV-2 are expected to change towards those of other coronaviruses that have been prevalent in humans and away from those of coronaviruses prevalent in bats. To test this hypothesis, we analyzed the following coronaviruses: 238 human-CoV strains (alphacoronaviruses 229E and NL63: betacoronaviruses HKU1 and OC43) and 166 bat-CoV strains (alphacoronaviruses and betacoronaviruses, including the SARS virus).

As shown in Fig. 3A, we compared the mononucleotide compositions of SARS-CoV-2 with those of the human- and bat-CoV strains; the data for bat SARS among bat-CoV strains, which is thought to be the original strain that caused the current COVID-19 pandemic, are marked in pink. Interestingly, concerning the human- and bat-CoV strains, differences in mononucleotide composition were more pronounced between hosts than between the alpha and beta linages, and the levels for all six clades of SARS-CoV-2 were between those for the two hosts. Fig. 3B shows the results of di- and trinucleotides, for which the directional, time-dependent changes were primarily common among the six clades. The increases and decreases in nucleotide composition observed for SARS-CoV-2 in Figs. 1 and 2 are indicated by hollow up and down arrows, respectively. Interestingly, all changes of interest tended to move away from the compositions of bat SARS and approach those of human-CoV, supporting the view that the directional changes of interest have biological significance and are possibly inevitable, as observed for influenza viruses. Assuming that approaching the levels in human-CoV strains is the hypothetical goal of the directional change of SARS-CoV-2, the current compositions are far from this hypothetical goal (Fig. 3); therefore, we predict that directional changes of interest will continue in the near future.

**Fig. 3.**
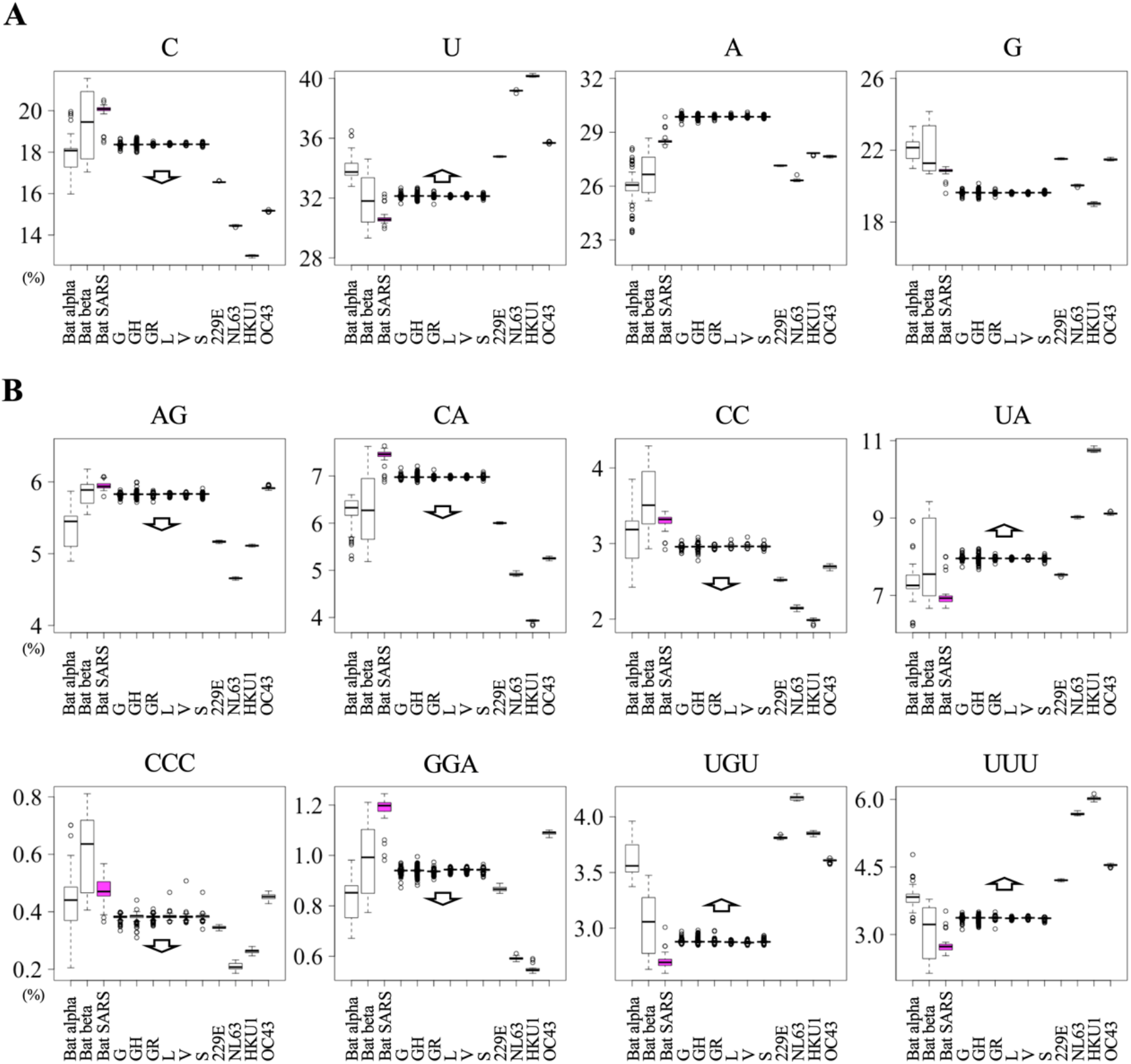
Nucleotide compositions of human and bat coronavirus sequences. A boxplot shows the nucleotide compositions in human-CoV (alpha 229E, alpha NL63, beta HKU1 and beta OC43), bat-CoV (bat SARS, alphacoronavirus and betacoronavirus) and SARS-CoV-2 strains. Bat SARS are marked pink. A hollow arrow indicates the direction of change in oligonucleotide composition observed for SARS-CoV-2 in Figs. 1 and 2. (A) Mononucleotides. To compare the four mononucleotides, the scale widths on the vertical axis scale are set to the same values. (B) Di- and trinucleotides.

Then, assuming that the average value for all human-CoV strains is a hypothetical goal, we investigated how SARS-CoV-2 has approached this possible goal. Specifically, we calculated the square of the difference between the composition of each nucleotide in SARS-CoV-2 and the average value for human-CoV strains and plotted the values of the difference according to the elapsed month for each nucleotide. Changes in the compositions of both C and U clearly reduced this difference, as the compositions of these nucleotides approached the hypothetical goal (Fig. 4A); their linear reduction supports the prediction that directional changes in the composition of C and U will continue for the foreseeable future. In contrast, A and G did not show directional changes in composition, which is most likely due to the absence of clear differences in the A and G compositions of human- and bat-CoV, i.e., there is no possible target (Fig. 3A). Fig. 4B shows examples of di- and trinucleotides whose compositions have moved towards the hypothetical goal, but Fig. 4C shows a few exceptional nucleotides whose compositions have not changed towards the hypothetical goal but have changed with a common directionality among the six clades. In Fig. 4D, correlation coefficients between the above difference and the elapsed month are presented. Most nucleotides of interest showed a negative coefficient (i.e., a directional change towards human-CoV), but three oligonucleotides, GG, AGC and CAU, showed positive coefficients indicating an increase in the difference (i.e., moving away from the human-CoV level). For these opposing directional changes, certain causes specific to SARS-CoV-2 may be assumed.

**Fig. 4.**
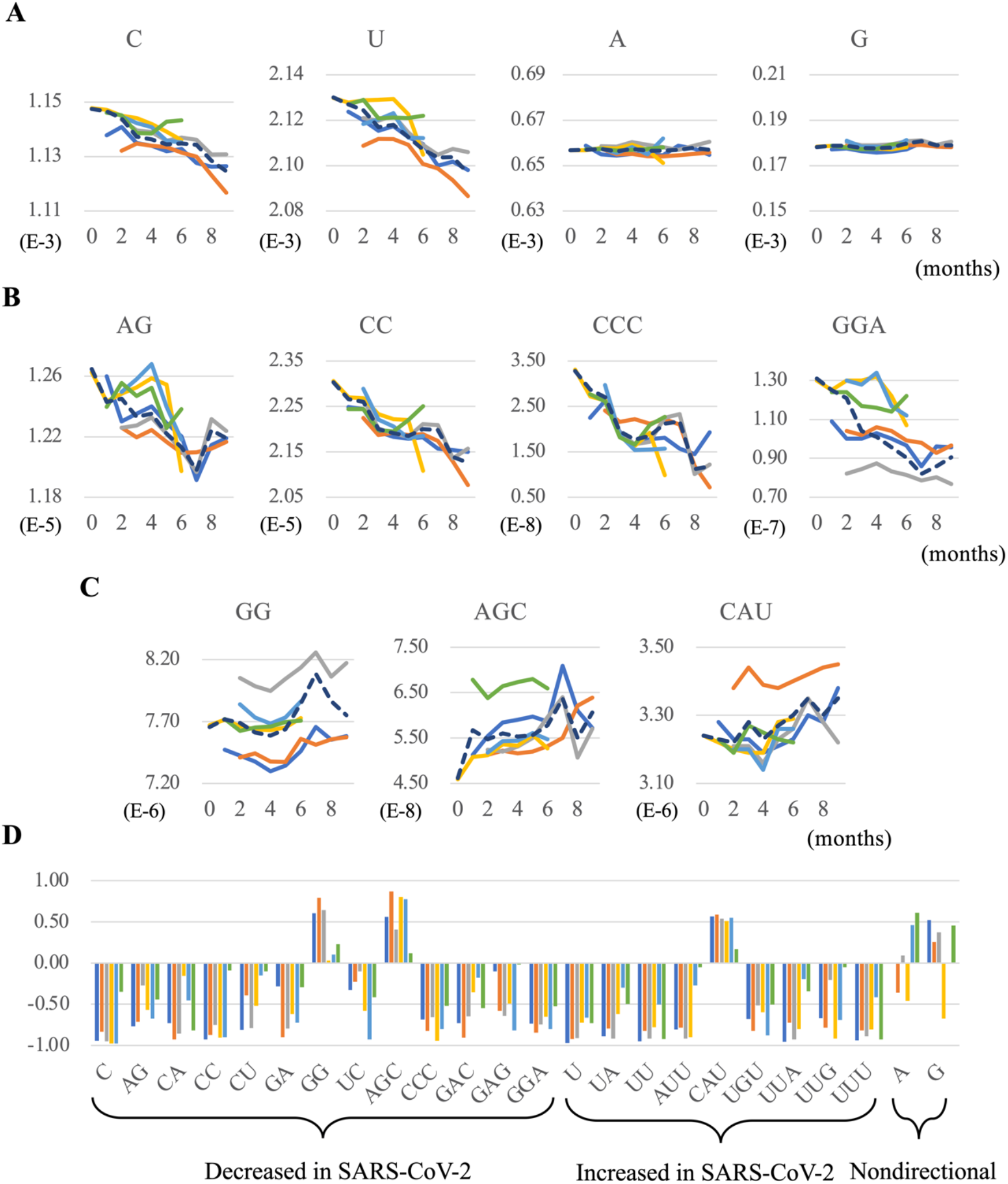
Differences in nucleotide composition between SARS-CoV-2 and human-CoV. (A) Values for the square of the difference in mononucleotide composition between SARS-CoV-2 isolated in each month and human-CoV are plotted against the elapsed month. The data are presented as colored or dashed lines, as described in Fig. 1. (B and C) Oligonucleotide compositions that approach and move from those of human-CoV are presented, respectively. (D) The correlation coefficients between the elapsed month from the start of the epidemic and the above differences in mono- and oligonucleotides whose directionality of change is common among six clades are presented. The results for A and G mononucleotides, which show nondirectional change, are also presented.

### Motifs for RNA-binding proteins

Next, we considered the mechanisms that move oligonucleotide compositions away from those of bat coronaviruses and closer to those of human coronaviruses. Certain human cellular factors involved in viral growth may be candidates in such mechanisms. When considering possible protein factors, oligonucleotides longer than trinucleotides should be a focus. As an attempt, we here focused on host RNA-binding proteins because their binding to hepatitis C virus is known to be involved in the growth of this RNA virus [20]. We thus searched for motifs for human RNA-binding proteins in coronavirus genomes (see Methods section) and found multiple loci with binding motifs for each protein. Table 3 (and Table S10) lists the motifs for which a directional time-dependent change was primarily common among six clades. Table 3 and Fig. 5A show that only ELAVL1 showed a positive correlation, but the other nine proteins in Table 3 showed a negative correlation for almost all clades; the results for other motifs are presented in Table S12.

**Table 3.** The number motif-containing loci for RNA-binding proteins whose occurrences have increased or decreased between strains of the first and last month of the analysis.

|  | Clade G | Clade GH | Clade GR | Clade L | Clade V | Clade S |
| --- | --- | --- | --- | --- | --- | --- |
| PTBP1 | -2.81 | -3.46 | -4.92 | -13.50 | -11.66 | 3.95 |
| HNRNPL | -3.03 | -1.78 | -0.52 | -3.62 | -6.48 | 0.71 |
| NOVA1 | -0.04 | -2.77 | -0.70 | -3.37 | -5.04 | 1.09 |
| SRSF2 | -1.17 | -3.02 | -1.09 | -3.10 | -3.10 | 0.78 |
| ZFP36 | 1.67 | -3.50 | -1.98 | -3.48 | -4.78 | 1.86 |
| HNRNPA1 | -0.16 | -2.50 | -0.50 | -2.64 | -3.93 | 0.44 |
| ELAVL1 | 2.82 | 0.47 | 2.87 | 0.59 | 0.03 | 2.39 |
| TIA1 | -0.82 | -2.05 | -1.20 | -1.72 | -4.41 | 1.67 |
| PCBP2 | -0.37 | -2.50 | -1.03 | -2.28 | -2.35 | 0.11 |
| SRSF1 | -0.63 | -2.29 | -1.26 | -2.55 | -1.84 | 0.36 |

**Fig. 5.**
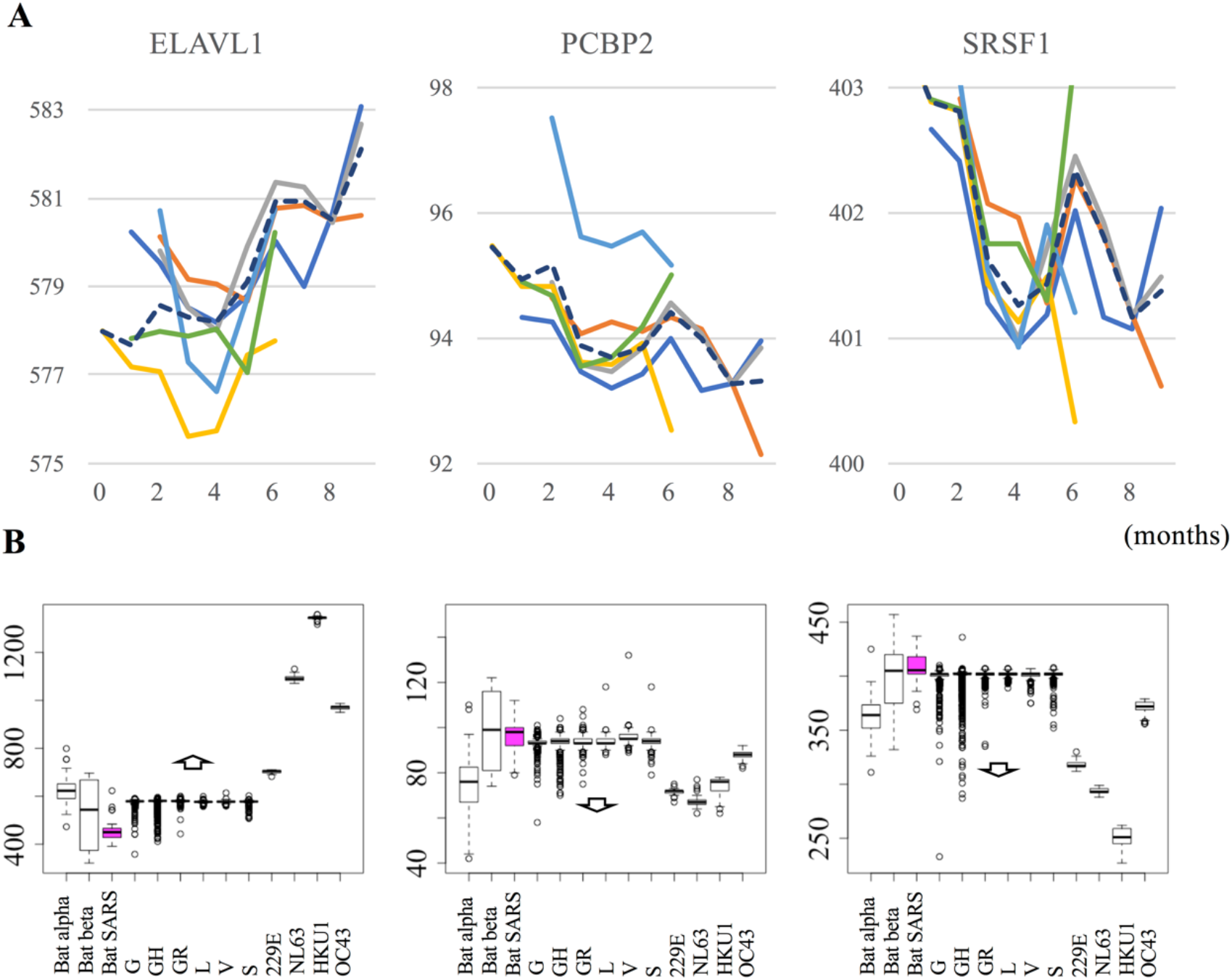
Time-dependent changes in the numbers of RNA-binding motif loci. (A) The numbers of loci containing RNA-binding motifs per genome are plotted against the elapsed month. Here, we selected RNA-binding proteins for which the number of motif loci increased or decreased by at least one for all six clades from the epidemic start. The data are presented as colored or dashed lines, as described in Fig. 1A. (B) A boxplot shows the number of loci containing RNA-binding motifs in human-CoV (alpha 229E and NL63: beta HKU1 and OC43), bat-CoV (bat SARS, alphacoronavirus and betacoronavirus) and SARS-CoV-2 strains. Bat SARS are marked pink. A hallow arrow indicates the direction shown in Fig. 5A with which the oligonucleotide compositions of SARS-CoV-2 changed.

We next compared the numbers of these motifs in SARS-CoV-2 with the numbers of human- and bat-CoV motifs (Fig. 5B). Of the ten proteins shown in Table 3, the only elevated motif, that for ELAVL1 binding, was found in a significantly higher number of loci in human-CoV than in bat-CoV, but motifs for PCBP2 and SRSF1 binding, which tended to decrease (Table 3), were found in significantly fewer loci in human-CoV. These observations appear to be consistent with the features found in the mono-, di- and trinucleotide compositions of interest. However, unlike these changes, there was significant diversity within even a single clade, which appears to be greater than the differences between hosts, with the possible exception of ELAVL1. In regard to long oligonucleotides, they should carry out a variety of functions, and mutations that accumulate in their functional motifs may have complex effects on the presence of functional motif sequences, so an analysis from a new perspective appears to become important.

## Discussion

We first discuss possible molecular mechanisms related to time-dependent directional changes in mononucleotide composition. Fig. 1A shows that the frequency of C tended to decrease in SARS-CoV-2, while that of U tended to increase. Since a similar change was previously found for MERS and all A-type influenza subtypes [12,14], these changes may have biological significance for a wide range of RNA viruses that invade from nonhuman hosts. One possible mechanism is the host RNA-editing function; Simmonds (2020) proposed that the C→U hypermutation in SARS-CoV-2 may be due to the influence of APOBEC family proteins in humans [19]. APOBEC is an antiviral protein in various animal species, including humans, that can convert C to U by the deacetylation of C [21–23]. Such RNA editing is also known to act as a defense mechanism against various viruses, including retroviruses [24]. The APOBEC gene family has generated various paralogs during mammalian evolution, with seven known APOBEC genes in humans and ten in bat families [25–27]. The prevalence C→U change in SARS-CoV-2 upon transfer of its host environment from bats to humans suggests that these changes may be due to human-specific APOBEC genes.

We next discuss changes in short oligonucleotides. Directional changes in some oligonucleotides, such as GAG and GGA, cannot be explained by APOBEC-induced C→U mutations alone. Although the evidence is weak, these oligonucleotides are part of the binding motifs of several RNA-binding proteins, such as SRSF1 and PCBP2 (Table S9); the number of loci for these motifs has decreased independently of clade. In contrast, the number of motif loci for only ELAVL1 among the ten proteins listed in Table 3 has increased independently of clade. As an RNA-binding protein that binds A- or U-rich elements, ELAVL1 binding to mRNA is known to contribute to RNA stability [28, 29]; SARS-CoV-2 and human-CoV, which are prevalent in humans, may contain increased binding motifs for ELAVL1 for efficient growth in the human cellular environment. However, for further analysis, information on RNA-binding proteins in bat cells is needed.

## Conclusions

In the present study, we found that the compositions of a group of mono- and oligonucleotide in SARS-CoV-2 genomes have changed in a host cell-dependent manner. This is totally consistent to our previous finding for influenza A and B viruses [11,12,14], supporting the previous prediction that the host-dependent directional changes of various mono- and oligonucleotides should inevitably occur in zoonotic RNA viruses that have invaded from nonhuman hosts. Phylogenetic methods based on sequence alignment [7,8] are well refined and undoubtedly essential for studying the phylogenetic relationships between viruses. The present alignment-free method to analyze mono- and oligonucleotide compositions can also serve as a powerful tool for molecular evolutionary studies of viruses, revealing directional changes in viruses and predicting the possible goals of these changes.

## Methods

### SARS-CoV-2 genome sequences

Human SARS-CoV-2 genome sequences were downloaded from the GISAID database (https://www.gisaid.org/); sequences that were complete, showed high coverage and had been isolated from humans were downloaded on Sep 17, 2020. Among the acquired sequences, strains with an unknown isolation month were excluded from the analysis, and the polyA tail was removed. A list of all 72,314 strains used is provided in Table S1.

### Genome sequences of coronaviruses prevalent in humans or bats

The complete sequences of two types of human coronavirus (human-CoV) strains, alphacoronaviruses (27 229E and 55 NL63 strains) and betacoronaviruses (18 HKU1 and 138 OC43 strains), were obtained from the NCBI virus database (https://www.ncbi.nlm.nih.gov/labs/virus/). The complete genome sequences of two types of bat coronavirus (bat-CoV) strains, alphacoronaviruses (87 strains) and betacoronaviruses (79 strains, including 34 SARS-CoV), isolated from three types of bats (Chiroptera, Vespertilionidae and Rhinolophidae) were obtained from the NCBI virus database (https://www.ncbi.nlm.nih.gov/labs/virus/), and the polyA tail of each sequence was removed. The strains are listed in Table S2.

### Time-series analysis of changes in oligonucleotide compositions

In the time-series analysis, the average mono- and oligonucleotide compositions (%) of viruses collected in each month were calculated for each clade. To avoid statistical fluctuations due to the small sample size, months in which fewer than 10 strains had been collected were excluded from the monthly analysis.

### RNA-binding motif analysis

RNA-binding motifs were obtained from the ATtRACT database [30]. In this database, multiple binding motifs are registered as corresponding to one RNA-binding protein; we calculated the total number of loci containing the binding motifs for each protein in the viral genomes.

## List of abbreviations

SARS-CoV-2: Severe acute respiratory syndrome coronavirus-2
human-CoV: human coronavirus
bat-CoV: bat coronavirus

## Ethics approval and consent to participate

Not applicable

## Consent for publication

Not applicable

## Availability of data and materials

The sequence dataset analyzed in this study are stored in GISAID. Other data are available from YI.

## Authors’ contributions

YI conceived the approach and conducted this analysis. TA developed the algorithm. TI supervised this study.

## Acknowledgements

We gratefully acknowledge the authors submitting their sequences from GISAID’s Database and also the valuable comments of Dr. Yashushi Hiromi of National Institute of Genetics (Mishima). We thank Springer Nature Author Services for editing this manuscript for English language.

## Additional file 1

Fig. S1: Average di- and trinucleotide compositions (A and B) of for SARS-CoV-2 strains collected in each elapsed month.

Fig. S2: Oligonucleotide compositions of human and bat coronavirus sequences.

Fig. S3: Differences in oligonucleotide composition between SARS-CoV-2 and human-CoV.

## Additional file 2

Table S1: List of SARS-CoV-2 strains used in the analysis.

Table S2: List of human-and bat-CoV strains used in the analysis.

Table S3: Number of SARS-CoV-2 strains in each clade isolated in each elapsed month.

Table S4: Average oligonucleotide compositions for SARS-CoV-2 strains in each clade isolated in each elapsed month.

Table S5: Correlation coefficients for time-dependent changes in oligonucleotide compositions of SARS-CoV-2.

Table S6: Fold change in compositions between strains of the first and last month of the analysis.

Table S7: Distance between the oligonucleotide composition of SARS-CoV-2 isolated in each elapsed month and that of human-CoV.

Table S8: Correlation coefficients for time-series changes in the distance between oligonucleotide compositions of SARS-CoV-2 and human-CoV.

Table S9: List of RNA-binding motifs.

Table S10: Numbers of motif-containing loci for RNA-binding proteins whose abundance increases or decreases between strains of the first and last month of the analysis.

Table S11: P-value from t-test to analyze the number of RNA-binding motif loci whose abundance increases or decreases between strains of the first and last month of the analysis.

Table S12: Correlation coefficients for time-dependent changes in the number of loci containing RNA-binding motifs.

