## Supplementary figures and images for "Human cell-dependent, directional, time-dependent changes in the mono- and oligonucleotide compositions of SARS-CoV-2 genomes"

### Additional_file_1_supply_fig

#### Slide 1
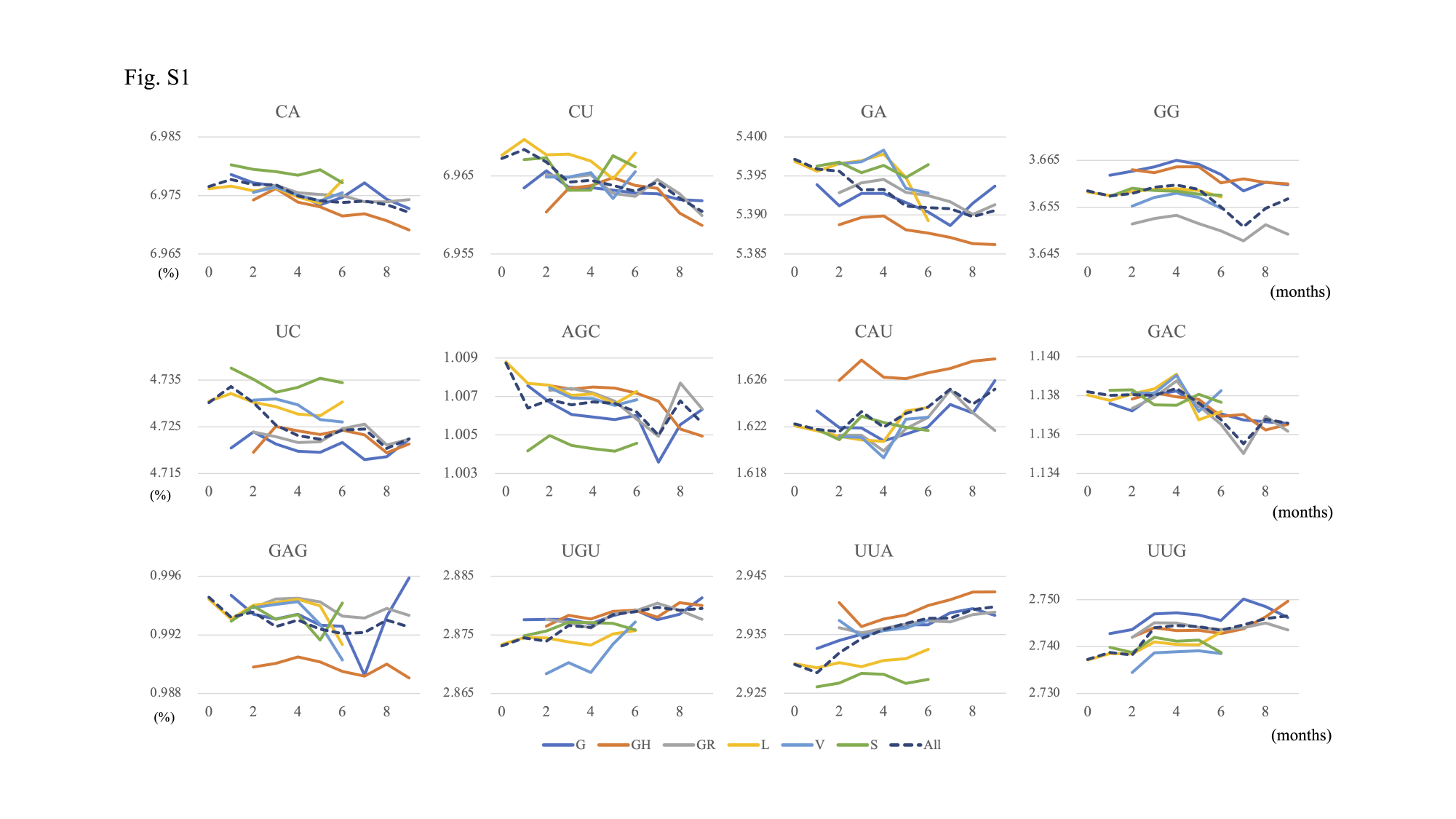

#### Slide 2
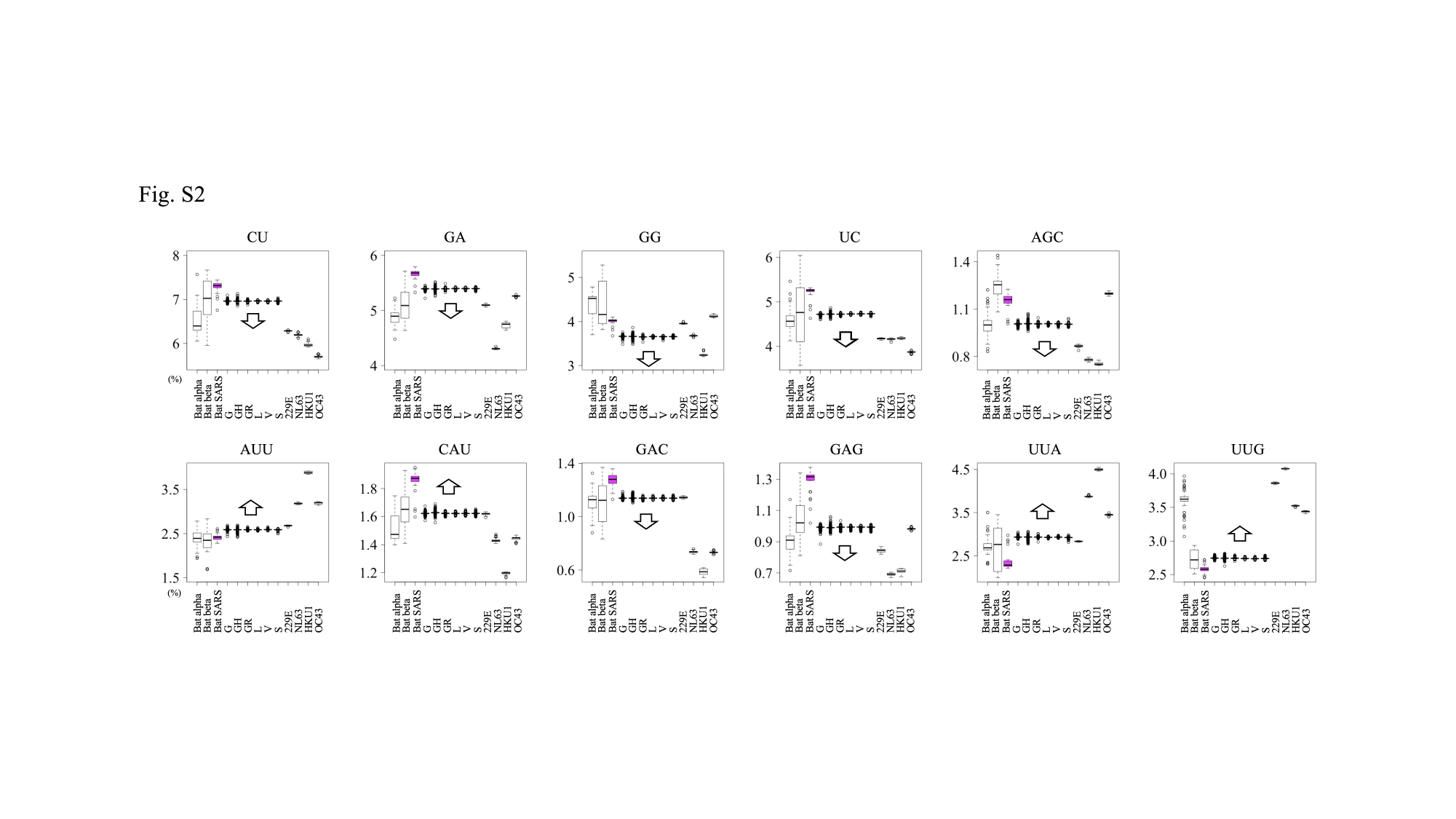

#### Slide 3
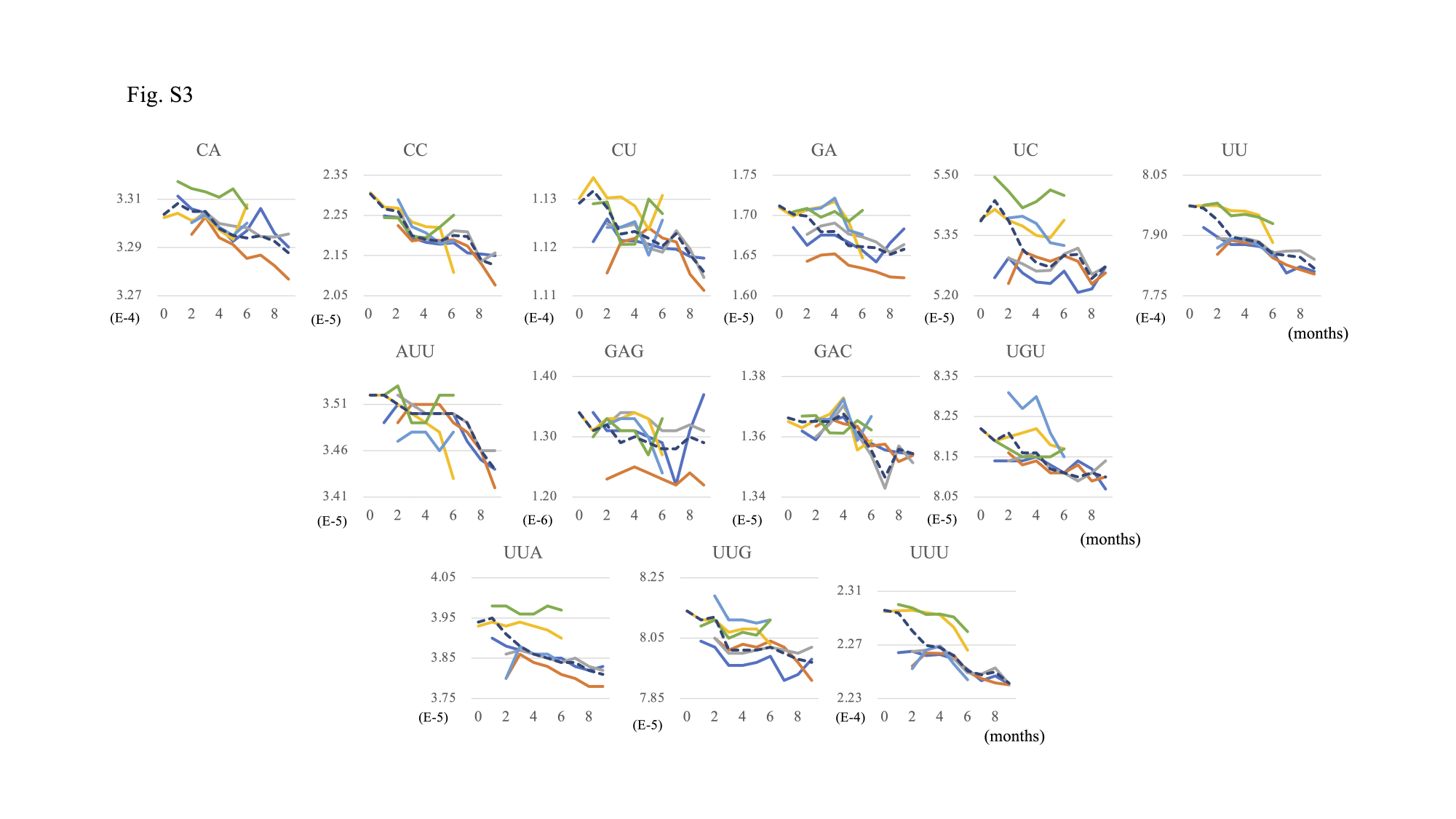
